# Using protein-per-mRNA differences among human tissues in codon optimization

**DOI:** 10.1101/2022.03.22.485268

**Authors:** Xavier Hernandez-Alias, Hannah Benisty, Luis Serrano, Martin H. Schaefer

## Abstract

Codon usage and nucleotide composition of coding sequences have profound effects on protein expression. However, while it is recognized that different tissues have distinct tRNA profiles and codon usages in their transcriptomes, the effect of tissue-specific codon optimality on protein synthesis remains elusive. Here, we leverage existing state-of-the-art transcriptomics and proteomics datasets from the GTEx project and the Human Protein Atlas to compute the protein-to-mRNA ratios of 36 human tissues. Using this as a proxy of translational efficiency, we build a machine learning model that identifies codons enriched or depleted in specific tissues. In particular, we detect two clusters of tissues with an opposite pattern of codon preferences. We then use the identified patterns for the development of CUSTOM, a codon optimizer algorithm which suggests a synonymous codon design in order to optimize protein production in a tissue-specific manner. In a human cell model, we provide evidence that codon optimization should indeed take into account particularities of the translational machinery of the tissues in which the target proteins are expressed and that our approach can design genes with tissue-optimized expression profiles. Altogether, CUSTOM could benefit biological and biotechnological research, such as the design of tissue-targeted therapies and vaccines.

## INTRODUCTION

From the advent of synthetic biology, it is widely recognized that gene design needs to be adapted to the expression requirements of the host^1^. Within coding sequences, there are manifold overlapping factors that determine translation, mRNA stability, transcription, splicing, methylation, or ribosomal frameshifting, among others^2^. Therefore, while the amino acid sequence of proteins is maintained, the usage of synonymous codons can be optimized for heterologous expression.

During the last decades, an extensive number of computational tools have been developed for gene design^3,4^. Most commonly, these tools optimize the codon usage in order to resemble that of the host based on the Codon Adaptation Index (CAI) of the genes to be optimized or similar metrics. Other more innovative developments also include neural networks that control translation speed^5^ or other machine learning algorithms that optimize mRNA stability^6^. Although there is no absolute “best” approach, codon optimization is commonly and successfully applied in gene design. In fact, current knowledge on the effect of synonymous variants on the heterologous expression of the protein GFP shows up to 46-fold expression differences in HeLa cells^7^. Similarly, mRNA and protein levels across thousands of GFP variants strongly correlated with their CAI in *S. cerevisiae*^8^.

Nevertheless, codon optimization in multicellular eukaryotes is more intricately determined, since different tissues can showcase differences in codon usage and tRNA expression^9–11^. The translational efficiency, which constitutes the rate of protein production from mRNA, is therefore dependent on the balance between the codon usage of genes being translated and the abundance of a limited tRNA pool^10,12^. In this context, codons translated by highly abundant tRNAs generally correspond to optimal codons in the translatome, as has been reported by ribosome profiling^13^. However, detecting differences of translational efficiency between tissues can be challenging, since the larger gene-to-gene variability of protein levels can obscure the actual tissue-to-tissue differences^14^.

The advent of high-throughput sequencing has enabled an extensive transcriptome profiling of human tissues^15,16^. Based on the mRNA-seq data from the GTEx project, Kames et al. (2020) developed the public resource TissueCoCoPUTs, containing codon and codon pair usage tables of tissue transcriptomes^11^. However, current knowledge indicates that tissue-specific variability of gene expression is mostly regulated at the post-transcriptional level and mRNA-seq alone is therefore not able to capture it^17,18^. Developments in mass spectrometry have very recently led to the release of deep and quantitative proteome maps of human tissues^19,20^.

Using this transcriptomic and proteomic data from the Human Protein Atlas and the GTEx project, we here compute the protein-to-mRNA (PTR) ratios of 36 human tissues as a proxy for translational efficiency. To distinguish high-PTR from low-PTR proteins, we build random forest models that identify which codons are optimal or non-optimal for each tissue. Then we apply these codon preferences to develop a tool, CUSTOM, that optimizes coding sequences for a specific tissue. CUSTOM is publicly available as a Python package (https://github.com/hexavier/CUSTOM) and as a web interface (https://custom.crg.eu). By optimizing eGFP and mCherry proteins to a human cell model of kidney and lung, we provide experimental evidence of how tissue codon optimization could be important e.g. in vaccines or gene therapy.

## RESULTS

### Protein-to-mRNA ratios detect differences in translational efficiency among tissues

Translational efficiency (TE) is defined as the rate of protein synthesis from mRNAs, which can be estimated as the protein-to-mRNA (PTR) ratio. To systematically analyze the PTR ratios across a total of 36 human tissues, we retrieved the mRNA-seq and proteomics data from two recent datasets: 29 tissues from the Human Protein Atlas^17,20^ (HPA) and 24 tissues from the GTEx project^19^ (**Figure 1A-B**, Supplementary Table 1). The first study includes one sample per tissue, which are concurrently analyzed by mRNA-seq and label-free iBAQ proteomics. On the latter, a total of 182 matched samples are measured both by mRNA-seq and tandem mass tag 10plex/MS3 mass spectrometry. By correlating the mRNA expression, protein abundance and PTR ratios along the 17 tissues in common, we could ascertain a high correspondence between the two datasets (ED Figure 1A).

**Figure 1.**
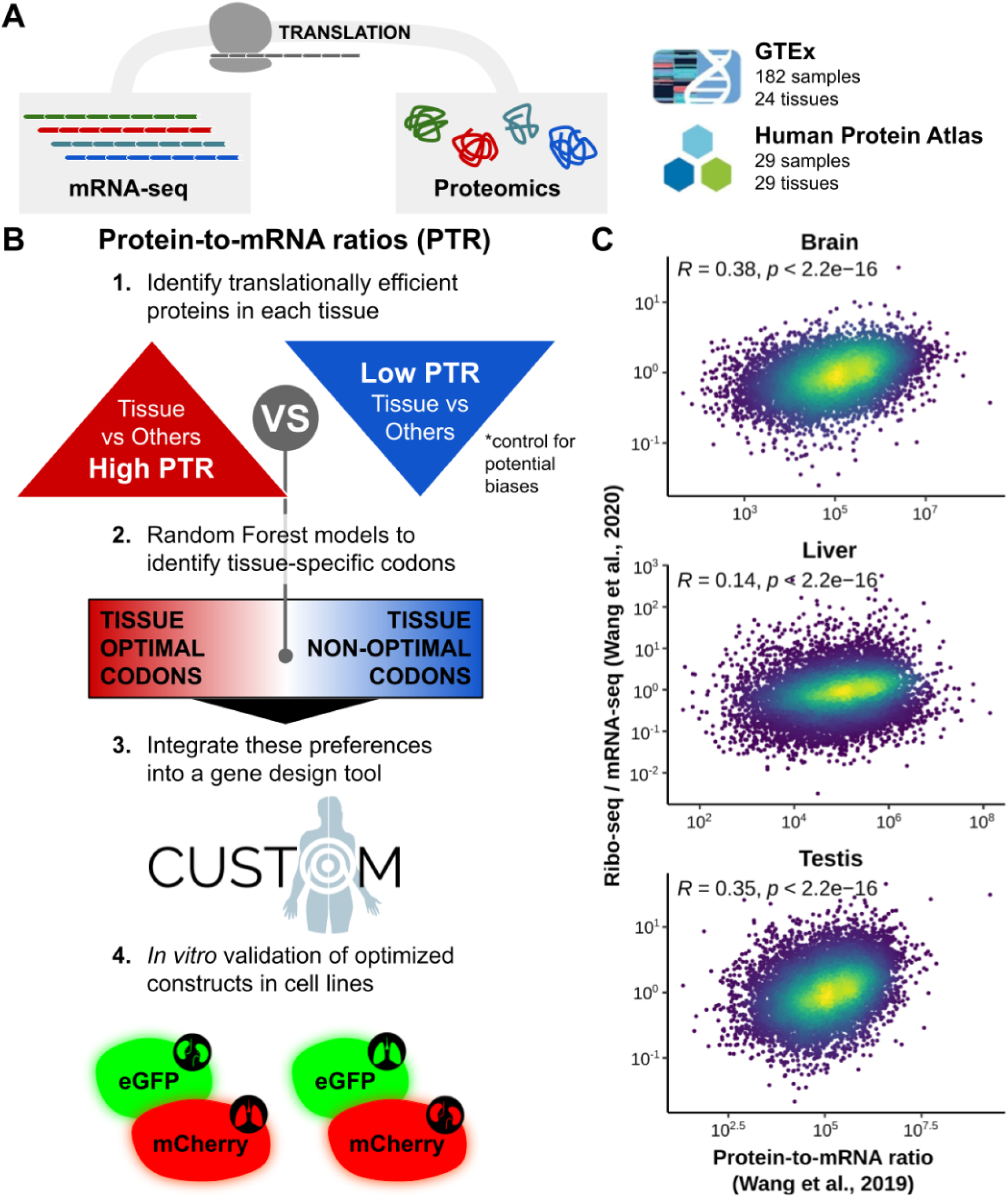
Protein-to-mRNA ratios detect differences in translational efficiency among tissues. (A) Proteomics and mRNA-seq data included in this study contains samples from the GTEx project^19^ and Human Protein Atlas^20^. (B) Using these datasets, we compute the protein-to-mRNA ratios (PTR) and define tissue-enriched and tissue-depleted sets of proteins for each tissue. By comparing the codon usage of these two sets, we identify the codon optimality pattern of tissues. Using this information, we develop a gene design tool called CUSTOM and validate the method using an *in vitro* cellular model. (C) Spearman correlation between the median translational efficiency^21^ (ratio between ribo-seq and mRNA-seq FPKMs) and PTR^20^ across genes in brain, liver, and testis. The color code depicts the density of points in the scatter plot.

Although to date this data is still relatively rare, a more direct readout of TE is the ratio between ribosome profiling and mRNA abundance. To confirm the validity of using PTR ratios as an estimate of TE, we therefore compared the PTR values to a ribosome profiling dataset of brain, liver, and testis. In all of them we observe a significantly positive correlation across the human genome^21^ (**Figure 1C**, Supplementary Table 1).

We next set out to investigate the tissue-to-tissue differences of PTR ratios in the aforementioned datasets. For each tissue, we defined a set of high-PTR and a set of low-PTR genes, described as having a PTR fold change compared to the average of all other tissues larger than 2, and vice versa (Supplementary Table 1). We find a significant concordance between the gene sets derived from the HPA and GTEx datasets in most tissues (p < 0.05, one-tailed binomial test, Supplementary Table 1).

To physiologically interpret the differences between gene sets, we performed an enrichment map among high-PTR and low-PTR sets linking tissues with high overlap of the respective gene sets (ED Figure 1B). In agreement with their highly tissue-specific function, we detect that tissues group according to their role in the body: eg. nervous tissue (brain and tibial nerve), muscular tissue (skeletal muscle and heart). Moreover, GO analyses of high-PTR genes show significant enrichments for highly tissue-specific biological processes according to the physiological and anatomical function of the tissue (p <0.05, Fisher’s exact test, ED Figure 1C).

We next asked if there could be any confounding factors associated with these gene sets, such as protein secretion and degradation, that could bias our analyses. On the one hand, it has been recently reported that constitutively secreted proteins are often detected at the mRNA but not at the protein level^19^, which could bias PTR ratios as a measure of TE. While we also observe these differences in our dataset (ED Figure 2A), the exclusion of secreted proteins from our gene sets does not affect the downstream results (see following section). On the other hand, we analyzed the protein half-life of gene sets based on two recent datasets in five human cell lines^22,23^ (Supplementary Table 1). The protein half-life is not significantly different between high-PTR and low-PTR gene sets in most of the tissues (p < 0.05, two-tailed Wilcoxon rank-sum test), nor is there any trend that one of the groups would be consistently associated with higher or lower half-life (ED Figure 2B).

Taken together, these observations indicate that PTR ratios can efficiently detect tissue-specific differences in translation. As such, it constitutes an appropriate dataset to systematically study TE differences across the set of 36 human tissues.

### Random Forest models identify two clusters of human tissues with distinct codon signatures

Recent studies show that different tissues can have different tRNA repertoires and codon usage^10,11^, which could have an influence on translational efficiency. Therefore, we wondered whether high-PTR and low-PTR sets of genes were specifically enriched or depleted of certain codons. If there is a tissue-specific codon signature, we would expect to be able to predict these differences in PTR.

To that aim, we built a random forest classifier for each tissue that predicts the high-PTR vs low-PTR state of genes based on their codon usage. All 36 resulting models perform with an area under the curve (AUC) of their receiver operating characteristic (ROC) curves higher than the no-skill model of 0.5 (**Figure 2A**, Supplementary Table 2). In particular, kidney, breast, lung, rectum and tonsil showcase the highest tissue-specific profiles (**Figure 2A**; all AUC > 0.70). Furthermore, to validate whether these differences in PTR are specifically dependent on codon usage and not from nucleotide composition alone, we compared them with the performance of three control models: +1 and +2 misframed codon usage as well as dinucleotide composition of genes (Supplementary Table 2). While these control models also show predictive power, the AUC of the correctly framed codon usage models significantly outperform the controls (p < 0.05, one-tailed binomial test).

**Figure 2.**
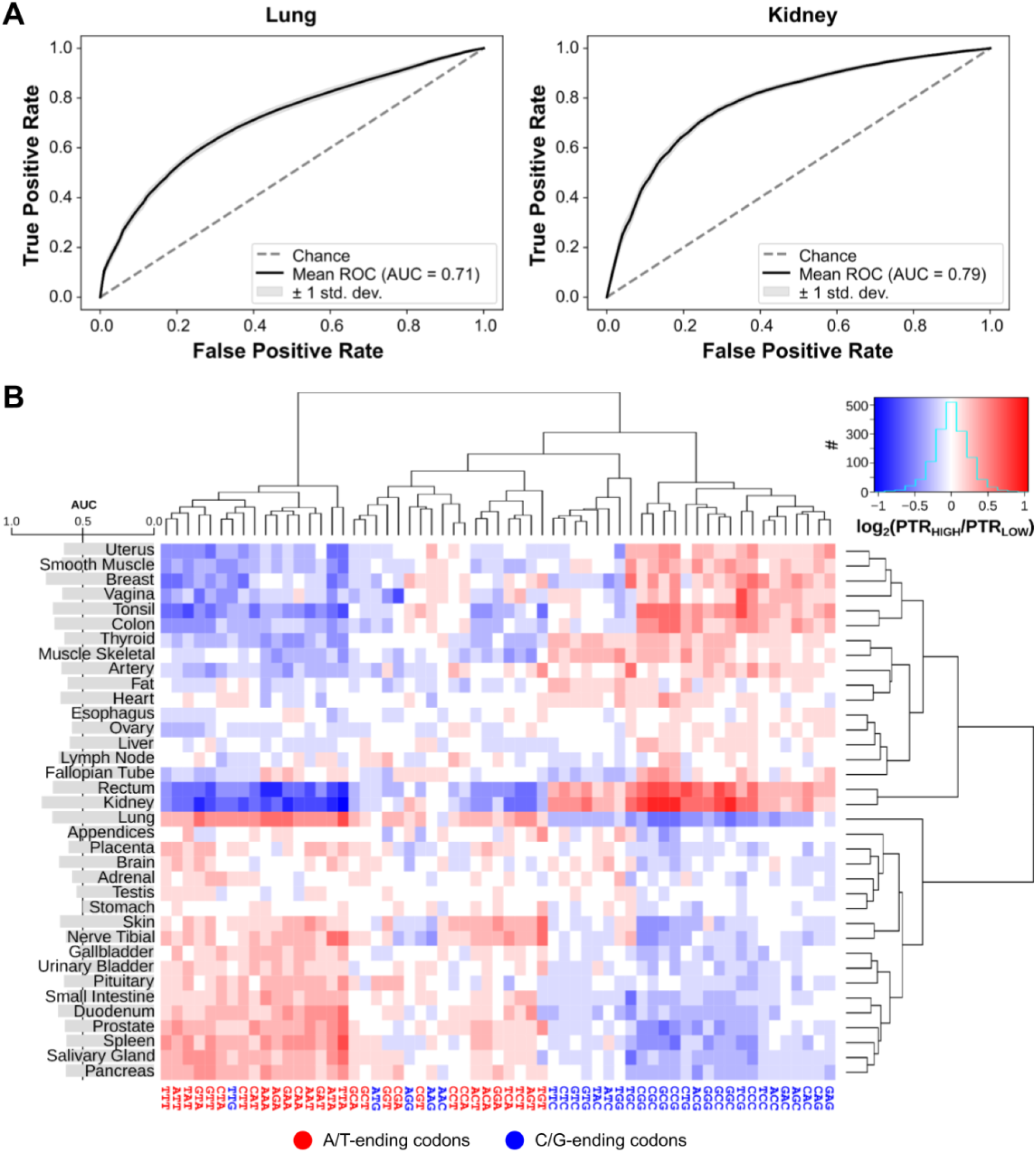
Random Forest models identify two clusters of human tissues with distinct codon signatures. (A) Receiver operating characteristic (ROC) curves of lung and kidney random forest classifiers, in which the codon usage of genes is used to predict whether they are high-PTR or low-PTR in the respective tissue (see Methods). (B) Ratios of the codon usage between high-PTR and low-PTR genes in each tissue. Codons and tissues are hierarchically clustered using euclidean distances and the complete-linkage method. The barplot on the left shows the mean AUC of the ROC curve of the RF model of each tissue.

To examine the tissue-specificity of codons, we next analyzed which particular codons are predictive for high vs low PTR states in each tissue. The relative feature importances of each random forest classifier measure the contribution of codons in the decision trees (ED Figure 3A). In general, only a few codons (5 to 10) are relevant for each model, but they differ across tissues. A recursive feature elimination of each model similarly substantiates that fewer than 10 codons are sufficient to achieve the maximum AUC performance (ED Figure 3B).

In addition, by computing the ratio between the codon usage of high-PTR vs low-PTR genes, we observe the enrichment or depletion of codons in specific tissues (**Figure 2B**). There are two main clusters of tissues with opposite codon optimality profiles: the first generally preferring A/T-ending codons while the second favoring C/G-ending ones. Also, as expected, tissues with higher AUC performances showcase more definite codon profile patterns both in terms of their enrichment/depletion (**Figure 2B**) as well as their importance (ED Figure 3A). As mentioned in the previous section, we also repeated the same analyses with the secretome-excluded sets of genes, which have a highly similar codon optimality profile with all correlations of codon ratios over 0.95 (Supplementary Table 2).

Given that some reports highlight the role of codon pair bias in translation^11,24^, we similarly analyzed the codon pair usage ratios between high-PTR vs low-PTR genes (Supplementary Table 3). A principal component analysis (PCA) of these ratios perfectly separates the exact same two clusters observed above with single codons alone (ED Figure 4A). To further analyze how much codon pair variance is explained by single codons alone, we compared observed codon pair ratios with their expected values based on their constituent single codons. They relate highly linearly as shown by linear regression models (ED Figure 4B, Supplementary Table 3), which indicates that differences in codon pair ratios can be explained by single codons alone. In fact, codon pairs that deviate the most from linearity just correspond to outliers with very low counts within gene sets (ED Figure 4C).

Overall, our random forest classifiers can predict the PTR of genes in a certain tissue based on their codon usage. As such, the observed differences in codon preference or avoidance across tissues can be exploited to optimize tissue-specific gene design.

### CUSTOM generates fluorescent variants with desired tissue-specific expression

To translate differences in tissue-specific PTR into a codon optimizer tool, we developed CUSTOM as a probabilistic approach (see Methods, https://custom.crg.eu). Given a certain amino acid sequence and a target tissue, codons are selected with a probability proportional to their tissue importance in the model (ED Figure 3A). Then, based on the ratio of the selected codon (**Figure 2B**), it is either added or avoided in the generated sequence. This process is performed along the whole sequence, and repeated iteratively to generate a pool of hundreds of optimized sequences. Among this pool of sequences, given that tissue-specific codon usage is not the only factor influencing coding sequences^2^, the top scoring ones can be selected based on other commonly used parameters of codon bias or mRNA stability^3^ (Codon Adaptation Index, Codon Pair Bias, Minimum Free Energy, Effective Number of Codons, see Methods).

To validate the predictor, we chose the proteins eGFP and mCherry, and optimized them with CUSTOM to either kidney or lung (Supplementary Table 4). Taking eight among the top optimized sequences (**Figure 3A**, 2x eGFP_Kidney_, 2x eGFP_Lung_, 2x mCherry_Kidney_, 2x mCherry_Lung_), we then designed four constructs, placing in each of them one eGFP and one mCherry optimized each one for a different tissue and under an inducible bidirectional promoter (**Figure 3B**). These constructs were then simultaneously expressed in the lung and kidney cell lines A549 and HEK293T, respectively. Based on available proteomics data of these cell lines^25^, the proteome of A549 clearly resembles that of lung, while HEK293 is a closer model to kidney (ED Figure 5A).

**Figure 3.**
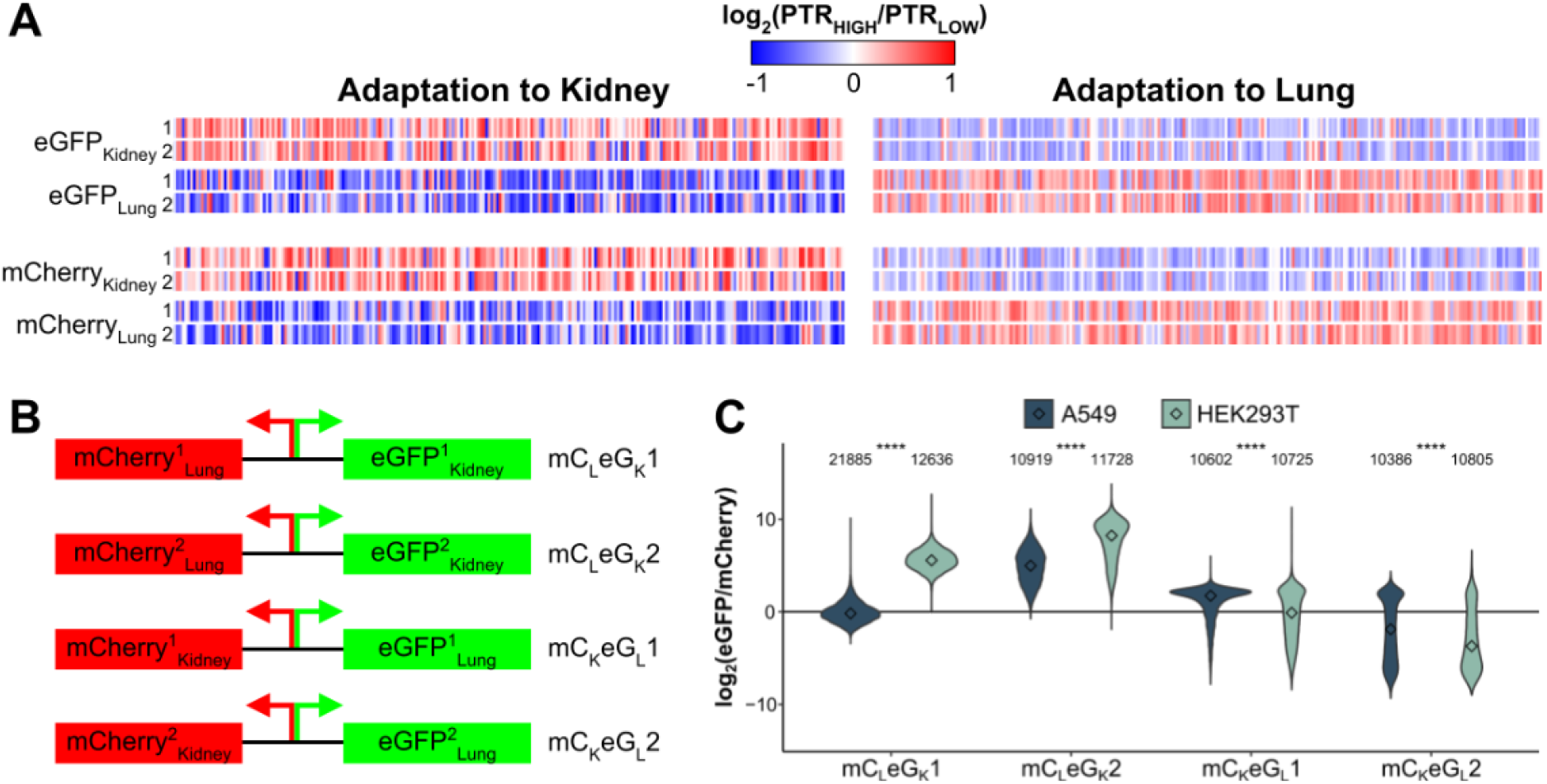
CUSTOM generates fluorescent variants with desired tissue-specific expression. (A) Selected eGFP and mCherry sequences optimized to lung and kidney using CUSTOM. The color code corresponds to the optimality ratios of Fig. 2B. (B) Using these sequences, we designed four of constructs by placing a mCherry and an eGFP with opposite tissue-specificity under an inducible bidirectional promoter. (C) Ratios of eGFP and mCherry for each of the four constructs detected by flow cytometry. The number of cells within each group is specified. Center values represent the median. Statistical differences were determined by two-tailed Wilcoxon rank-sum test, and are denoted as follows: *p ≤ 0.05, **p ≤ 0.01, ***p ≤ 0.001, ****p ≤ 0.0001. Two additional replicates are shown in ED Figure 5A.

We then analyzed the eGFP and mCherry fluorescence of each construct in each cell line. For all cases, we observe that the eGFP/mCherry ratio is significantly higher in the tissue for which eGFP is optimized (**Figure 3C**, p < 0.05, two-tailed Wilcoxon rank-sum test, ED Figure 5B), which validates our tissue-specificity hypothesis. We further observe that (1) the two constructs with eGFP_Lung_ have generally lower eGFP/mCherry ratios compared to the ones with eGFP_Kidney_, and (2) the differences in eGFP/mCherry ratios between constructs are more variable in HEK293 than A549 cells. Altogether, these observations suggest that A/T-ending codons are generally lower expressed than C/G-ending counterparts, but tissues like lung tolerate them better.

## DISCUSSION

Current analyses of the mRNA and protein levels among human tissues distinguish between across-gene and within-gene (i.e. across-tissue) variability^14^. In fact, the coefficient of variation of mRNA and protein levels across genes highly exceeds that of across tissues. In consequence, studies of codon usage on human transcriptomes and PTR ratios so far were dominated by the across-gene variability, and thus overlooked the smaller across-tissue differences^11,17^. The approach taken here puts the focus on the across-tissue variability of PTR ratios rather than the overall genome, which is actually the major source of post-transcriptional regulation^17,18^. In fact, we provide evidence that high-PTR gene sets of tissues are particularly enriched for tissue-specific functions.

Given the high GC content of the human genome as a whole, G/C-ending codons are generally more abundant (i.e. higher CAI), and relate to higher mRNA and protein expression levels^7,26,27^. But again, moving away from this across-gene perspective of human codon usage to look at the across-tissue variation, we here report that distinct tissues showcase different codon preferences. All in all, as also determined experimentally, we observe that the expression of a certain protein is dependent on two axes: (1) the across-gene axis with G/C-ending codons favoring higher absolute expression, and (2) the tissue-specific axis with the codon preferences observed in Figure 2B. Moreover, we also report that some tissues have a more definite codon profile than others, where this second axis is less evident. In agreement with our observed tissue-specific axis, Allen et al. (2022) recently reported that testis and brain (in contrast to other tissues such as ovary) better tolerate the translation of rare A/T-ending codons in *Drosophila melanogaster*^28^.

The codon optimization tool CUSTOM is able to exploit these codon preferences for the design of tissue-targeted genes. In fact, all four designed constructs expressed in a kidney and lung cell line showed the predicted tissue-specificity. To make CUSTOM readily available to the community, we developed it completely open source and made it accessible through a web server.

Human tissues are ensembles of heterogeneous cell types, and therefore observed differences in codon optimality are actually a composition of the constituent cell types. However, single-cell technologies of mRNA and protein measurements fall still far from complete cellular atlases^29^. Instead, we used the most up-to-date and complete tissue-wide maps of the human transcriptome and proteome, which have been generated by cutting-edge mass spectrometry and mRNA sequencing techniques^15,19,20^.

Finally, the results presented here constitute a proof-of-concept that tissue-specific codon usage exists and can be applied to gene design. In particular, this tool could be used in the development of optimized gene therapies or mRNA vaccines with more targeted tissue targets and therefore potentially less side effects. Nevertheless, factors other than codon usage also play a role in gene expression^2^, and therefore changes in synonymous codons can as well interfere with other processes such as mRNA folding and stability, mRNA modifications, protein folding, or translational kinetics^30,31^. As such, tissue-specific codon usage will constitute one additional instrument in the gene design tool set.

## METHODS

### Codon optimizer for tissue-specific expression

CUSTOM is implemented in Python (version >= 3.7) and available on GitHub (https://github.com/hexavier/CUSTOM) and as a web interface (https://custom.crg.eu). The landscape of possible synonymous sequences is vast and manifold factors overlap in defining the code. Therefore, we follow a simple probabilistic approach with two steps: (1) translate tissue-specific codon preferences into a pool of optimal sequences, and (2) select the desired sequence based on other parameters of relevance.

#### Create a pool of tissue-optimized sequences

The algorithm requires two main input data: the amino acid sequence to be optimized (or DNA sequence) and the target tissue. For each iteration of the optimization, the sequence is optimized taking two factors into account: how important the codon is in defining tissue-specificity (relative feature weights in ED Figure 3A) and whether it is enriched or depleted in the tissue (codon ratios in Figure 2B). Therefore, for each amino acid, a certain codon is selected with a probability proportional to the first. If the selected codon is enriched in the tissue, it is incorporated into the sequence. If it is depleted, the codon is excluded and another codon is selected based on the same probabilities as before. This process is repeated along the full sequence, and for as many iterations as desired. Furthermore, given that 5-10 top codons are often sufficient to achieve the full AUC prediction (ED Figure 3B), users can also control whether optimizing all codons or only the top ones.

#### Selecting the top scoring candidates

Once a pool of optimized sequences has been generated, the best-ranked ones can be selected as the user desires. Given that no ground truth is known, the default *select_best* method of the package measures a list of standard metrics frequently used in gene design and computes an average to select the top scoring sequences. The following factors can be included:

- Minimum Free Energy (MFE): a measure of mRNA stability from the ViennaRNA package^32^. CUSTOM distinguishes between the first 40 nucleotides (whose weak secondary structure leads to increased translation initiation) and the rest of the sequence (whose strong secondary structure relates to longer mRNA half-lives)^3^.
- Codon Adaptation Index (CAI): a measure of similarity between the codon usage of the sequence and that of the human genome^33^.
- Codon Pair Bias (CPB): a measure of similarity between the codon pair usage of the sequence and that of the human genome^24^.
- Effective Number of Codons (ENC): a measure of codon evenness. A value of 20 means that all 100% codons are biased towards the most common codon, while 61 corresponds to no bias at all^34^.
- GC content: a measure of similarity between the sequence GC content and a desired target value of GC.
- Homopolymers: filters out sequences with homopolymers of a certain length, which can lead to worse expression.
- Motifs: filters out sequences containing certain motifs.

### Experimental model and protocol

#### Human cell models

The cell lines included in this study are HEK293T and A549. The sex of each cell line is as follows: HEK293T, female; A549, male. Cells were maintained at 37°C in a humidified atmosphere at 5% CO2 in DMEM 4.5 g/l Glucose with UltraGlutamine media supplemented with 10% of FBS and 1% penicillin/streptomycin.

#### Expression vectors design

We applied CUSTOM to the protein sequences of eGFP and mCherry (Uniprot ID: C5MKY7, X5DSL3). Sequences were optimized to either lung or kidney, generating a total of *n_pool* = 1000. Sequences with homopolymers equal or larger than 7 were filtered out and scored with:

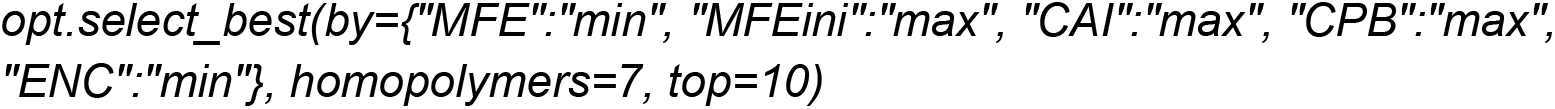

Among the top 10 scoring candidates of each optimization, we selected 2x eGFP_Kidney_, 2x eGFP_Lung_, 2x mCherry_Kidney_, and 2x mCherry_Lung_ (Supplementary Table 4).

For gene overexpression experiments, the two selected eGFP and and mCherry were cloned into a modified version of the XLone-GFP vector (Addgene#96930). The modification consisted of replacing the promoter of XLone-GFP with a bidirectional TRE3G promoter (Clontech), which allows the simultaneous expression of both genes. The four constructs consisted in a combination of eGFP_Lung_ + mCherry_Kidney_ and eGFP_Kidney_ + mCherry_Lung_.

#### Flow cytometry

HEK293T and A549 cells were seeded in 6-well plates. Gene expression was induced with 500 ng/mL of doxycycline during 48h. To measure the expression of the fluorescent proteins, cells were trypsinized and resuspended with 500 μL of media. Samples were applied on a FACS Fortessa analyser. Approximately 10^4^ live single-cell events were collected per sample. BD FACSDiva software was used for gating and analysis. The fluorescence intensity for each population in the FITC channel and PE–Texas Red channel was obtained.

### Data sources

#### Protein-to-mRNA ratios

The PTR ratios of the HPA were directly retrieved from the Table EV3 of Eraslan et al. (2019). In this dataset, protein levels are determined as absolute abundances based on their iBAQ quantification. As for the GTEx data, we retrieved protein and mRNA levels from Table S2 and Table S3 of Jiang et al. (2020), respectively. In this case, the proteomics measurements are relative quantifications from a tandem mass tag (TMT) 10plex/MS3 mass spectrometry strategy. To compute their PTR ratios, we followed the same pipeline as in the HPA: (1) proteins with an abundance of 0 were considered as missing values (NA); (2) protein quantifications were adjusted to have in each tissue the same median than the overall median; (3) genes with a TPM lower than 10 were taken as non-transcribed (NA). With that, comparable PTR values between HPA and GTEx are obtained (ED Figure 1A).

#### Codon and codon pair usage tables

The codon usage and codon pair usage tables of *Homo sapiens* from RefSeq were downloaded from the Codon/Codon Pair Usage Tables (CoCoPUTs) project release as of June 9^th^, 2020^35^. Regarding the codon usage of misframed coding sequences and their dinucleotide composition, we computed them from the latest release of the CCDS database of human sequences (release 22)^36^.

#### Translational efficiencies

The processed data of matched ribosome profiling and mRNA-seq samples from brain, liver and testis was retrieved from ArrayExpress (E-MTAB-7247)^21^. Translational efficiencies were then computed as the ratio FPKM_Ribo-seq_/FPKM_mRNA-seq_.

#### Protein half-life

The log-10-transformed protein half-lives for B cells, NK cells, hepatocytes, monocytes, and HeLa cells were downloaded from Eraslan et al. (2019)^22,23^. Given the concordance of half-lives among the five cell types (Supplementary Table 1), we used their average for the analysis in this work (ED Figure 2B).

#### Blood secretome

Using the predictions by the HPA^16^, there are 2641 secretome genes, 729 of which are secreted to blood. Given that we were concerned on proteins that are not detected at the protein levels because of their systemic rather than local secretion, we focused our analysis on the latter (Supplementary Table 1).

### Computational analysis

#### High-PTR and low-PTR gene sets

As PTR values from GTEx were computed from relative TMT proteomics in contrast to the absolute iBAQ quantification of HPA, they were not directly comparable and thus we defined the high-PTR and low-PTR gene sets for each dataset separately. On the one hand, high-PTR genes fulfilled three conditions: (1) genes having a PTR fold change compared to the average of all other tissues larger than 2, (2) genes with the highest PTR among all tissues, (3) genes detected in at least 3 tissues in the dataset. On the other hand, low-PTR genes were defined as: (1) genes having a PTR fold change compared to the average of all other tissues smaller than 0.5, (2) genes with the lowest PTR among all tissues, (3) genes detected in at least 3 tissues in the dataset. As a result, we defined one high-PTR and one low-PTR gene set for each tissue in each dataset. For those 17 tissues in common between both HPA and GTEx datasets, the union between both datasets was taken except for genes with contradictory labels, which were excluded.

#### Random Forest classifiers

To identify the most important codons determining high-PTR vs low-PTR genes, we computed their codon usage normalized by length, so that all 61 amino-acid-encoding codons sum up to 1. Taking this table of normalized codon usage as features, we applied a Random Forest (RF) classifier, populated with 100 decision trees, using the scikit-learn package^37^. Therefore, for each of the 36 tissues, we developed a model for predicting the high-PTR vs low-PTR genes based on their codon usage. To control for size differences between high-PTR and low-PTR groups, we iteratively sampled equal-sized groups, for n = 100 iterations. Furthermore, we validated the results with a stratified 5-fold cross-validation. In order to evaluate the performance of the RF models, we computed the Area Under the Curve (AUC) of Receiver Operating Characteristic (ROC) plots (Figure 2A). We took the average and standard deviation across all iterations. Similarly, we computed the relative feature weights corresponding to each of the 61 codons (Figure 2B).

To validate that the predictive potential of RF classifiers were codon-specific, we similarly computed the length-normalized codon usage of +1 and +2 misframed coding sequences as well as dinucleotide usage. By running the exact same pipeline as above, we determined the average AUC of these three control RF classifiers (Supplementary Table 2). We used a one-tailed binomial test to analyze whether the AUCs of controls were lower than the original model more often than expected by chance (p = 1/2).

While the relative feature weights determine the importance of each codon in distinguishing high-PTR vs low-PTR genes, they do not provide any directionality. To analyze whether codons are enriched or depleted in high-PTR vs low-PTR genes, we computed the ratios between the average length-normalized codon usage of high-PTR and low-PTR genes. Similarly, codon pair ratios were computed in the same way.

Among the total of amino-acid-encoding 61 codons, we also analyzed how many of them were actually informative in the models using a Recursive Feature Elimination (RFE). Therefore, for each tissue, we started by building a full model with all 61 codons and then recursively removed the least important one, as determined by the relative feature weights, until only one was left. At each step, we computed the AUC of the ROC curve of the model as explained above (ED Figure 3B).

#### Enrichment map

For this analysis, in order to allow an overlap between tissue gene sets, we used a slightly less stringent tissue-specificity definition. High-PTR sets were defined as (1) genes having a PTR fold change compared to the average of all other tissues larger than 2 and (2) genes detected in at least 3 tissues in the dataset, and vice versa for low-PTR sets.

To analyze the overlap between tissue gene sets, we used the EnrichmentMap app from Cytoscape^38^. We defined a generic input of high-PTR and low-PTR sets of proteins per tissue. Similarity was computed as the overlap coefficient ([size of (A intersect B)] / [size of (minimum(A, B))]).

#### Gene Ontology enrichment analysis

Gene Ontology (GO) categories of Biological Processes were analyzed for enrichment as of May 27^th^, 2021^39^. Enrichment analyses were performed by PANTHER using the Fisher’s exact test and Bonferroni correction for multiple testing^40^.

#### Principal Component Analysis of codon pairs

We applied Principal Component Analysis to the codon pair ratios of each tissue in order to explore the main variability among tissues along the 4096 codon pair ratios.

#### Linear regression of codon pairs

We fitted a linear regression model between the observed codon ratios (dependent variable) and the expected ratios based on single codons alone (independent variable). The expected values were computed as the product of the ratios of the two codons that constitute the pair. For each model, we computed the R squared, the Residual Standard Error (RSE), and the model p-value (Supplementary Table 3).

#### Statistical analysis

All details of the statistical analyses can be found in the Results section and the figure legends. We used a significance value of 0.05.

## ACKNOWLEDGEMENTS

We acknowledge the support of the Spanish Ministry of Science and Innovation (MICINN) (PGC2018-101271-B-I00 Plan Estatal), ‘Centro de Excelencia Severo Ochoa’, the CERCA Programme/Generalitat de Catalunya. The work of X.H. has been supported by a PhD fellowship from the Fundación Ramón Areces.

## AUTHOR CONTRIBUTIONS

Conceptualization, X.H., H.B., M.H.S. and L.S.; Methodology, X.H., H.B., M.H.S., L.S.; Software, X.H.; Investigation, X.H., H.B.; Validation, X.H., H.B., M.H.S.; Formal analysis, X.H., H.B.; Writing-Original Draft, X.H., H.B.; Writing-Review & Editing, X.H., H.B., M.H.S., L.S.; Visualisation: X.H., H.B., M.H.S., L.S.; Funding Acquisition, L.S.; Supervision, M.H.S. and L.S.

## DATA AND CODE AVAILABILITY

The code used in this study is available at GitHub (https://github.com/hexavier/codon_optimization), and the CUSTOM software is available as a Python package (https://github.com/hexavier/CUSTOM) and a web interface (https://custom.crg.eu). The published article includes all datasets generated or analyzed during this study.

## EXTENDED DATA FIGURES

**Extended Data Figure 1.**
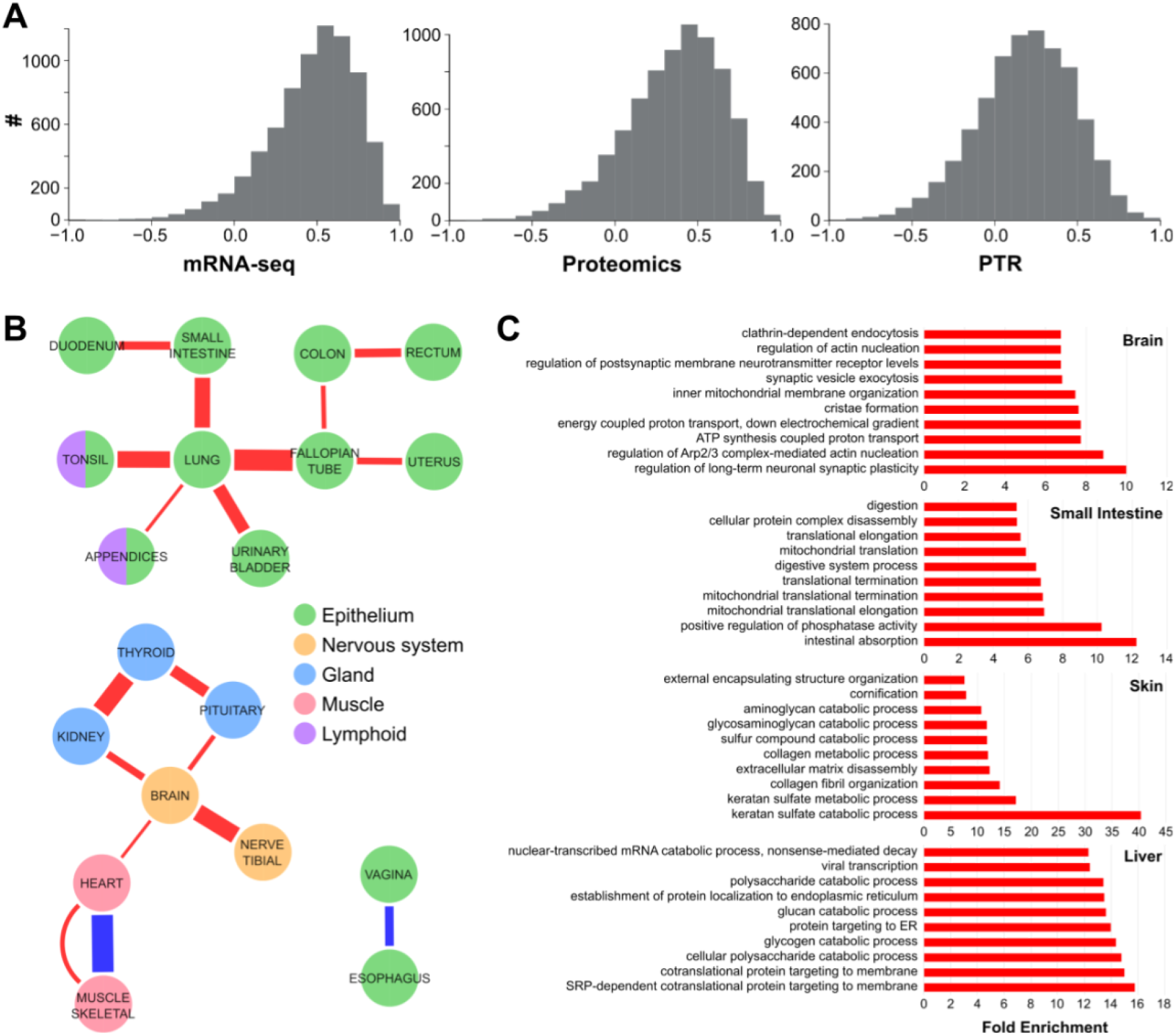
Protein-to-mRNA ratios detect differences in translational efficiency among tissues, related to Figure 1. (A) Correspondence between GTEx and Human Protein Atlas datasets at the mRNA-seq, proteomics and PTR levels. The histograms show the Spearman correlation of each gene along all 17 tissues in common between both datasets. Only correlations of genes detected in more than 5 tissues were computed. (B) Enrichment Map of high-PTR (red) and low-PTR (blue) sets of proteins among tissues. Edges show significant enrichments between sets with a similarity coefficient >0.33. The width of edges is proportional to the similarity coefficient. The size of nodes is proportional to the number of genes in the set, and their color depicts their tissue type based on the BRENDA Tissue Ontology^41^. Tissues with no significant edges are not shown. (C) GO enrichment analysis of biological processes for the high-PTR sets of four tissues. The top 10 significant GO terms with a Bonferroni-corrected p-value ≤ 0.05 are shown.

**Extended Data Figure 2.**
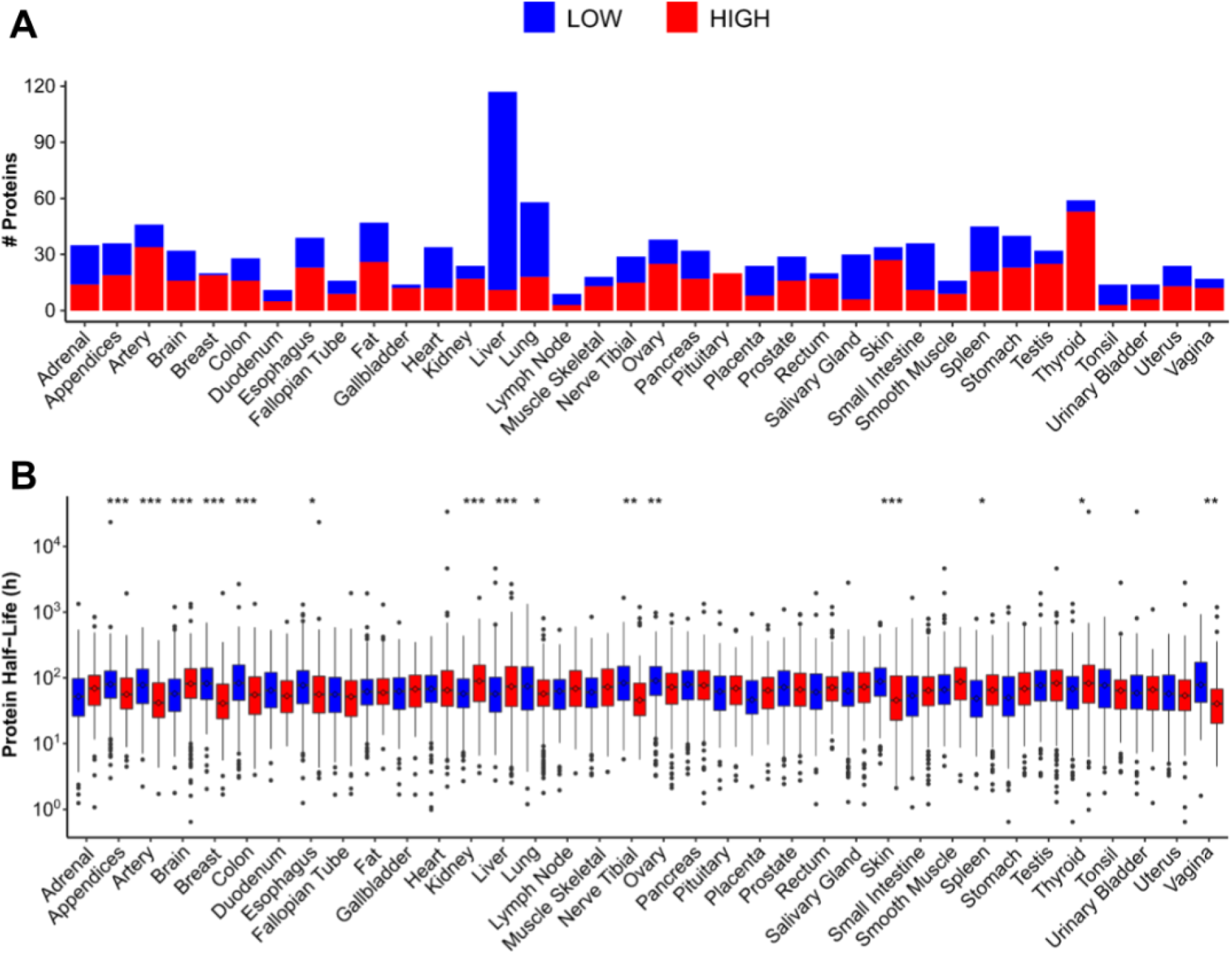
Differences in secretion and protein half-life among tissues, related to Figure 1. (A) Number of proteins secreted to blood in the low-PTR and high-PTR sets of genes in each tissue. (B) Average protein half-life of low-PTR and high-PTR sets of genes per tissue. Statistical differences were determined by two-tailed Wilcoxon rank-sum test and corrected for multiple comparisons using the Holm-Bonferroni method. Only significant differences are shown and are denoted as follows: *p ≤ 0.05, **p ≤ 0.01, ***p ≤ 0.001.

**Extended Data Figure 3.**
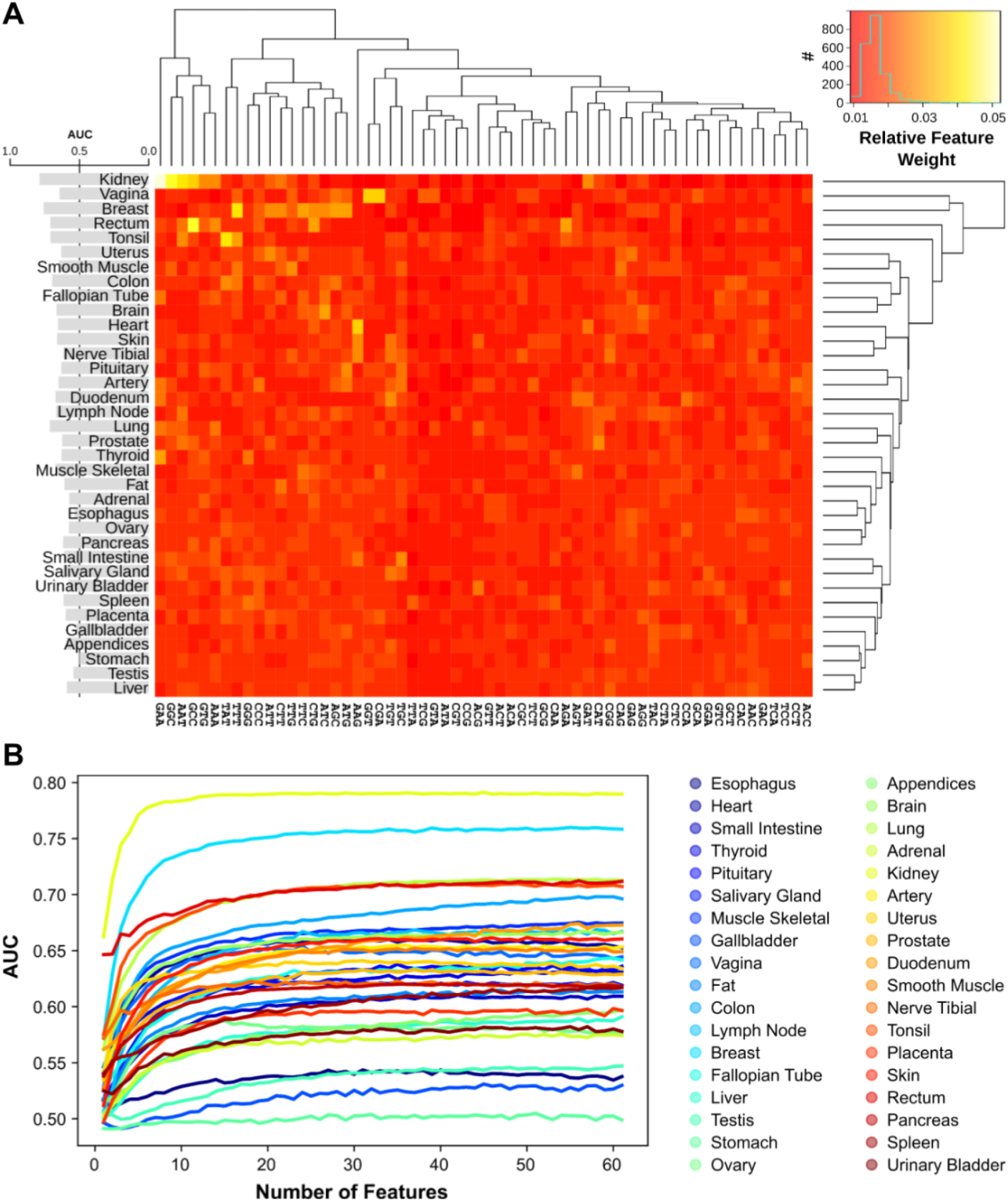
Random Forest models identify two clusters of human tissues with distinct codon signatures, related to Figure 2. (A) Based on the random forest classifiers, the heatmap shows the relative feature weights of every codon for each of the 36 human tissues, which measure the contribution of each codon in the decision trees. Codons and tissues were hierarchically clustered using euclidean distances and the complete-linkage method. The barplot on the left shows the mean AUC of the ROC curve of the RF model of each tissue. (B) Recursive Feature Elimination of the random forest model of each tissue.

**Extended Data Figure 4.**
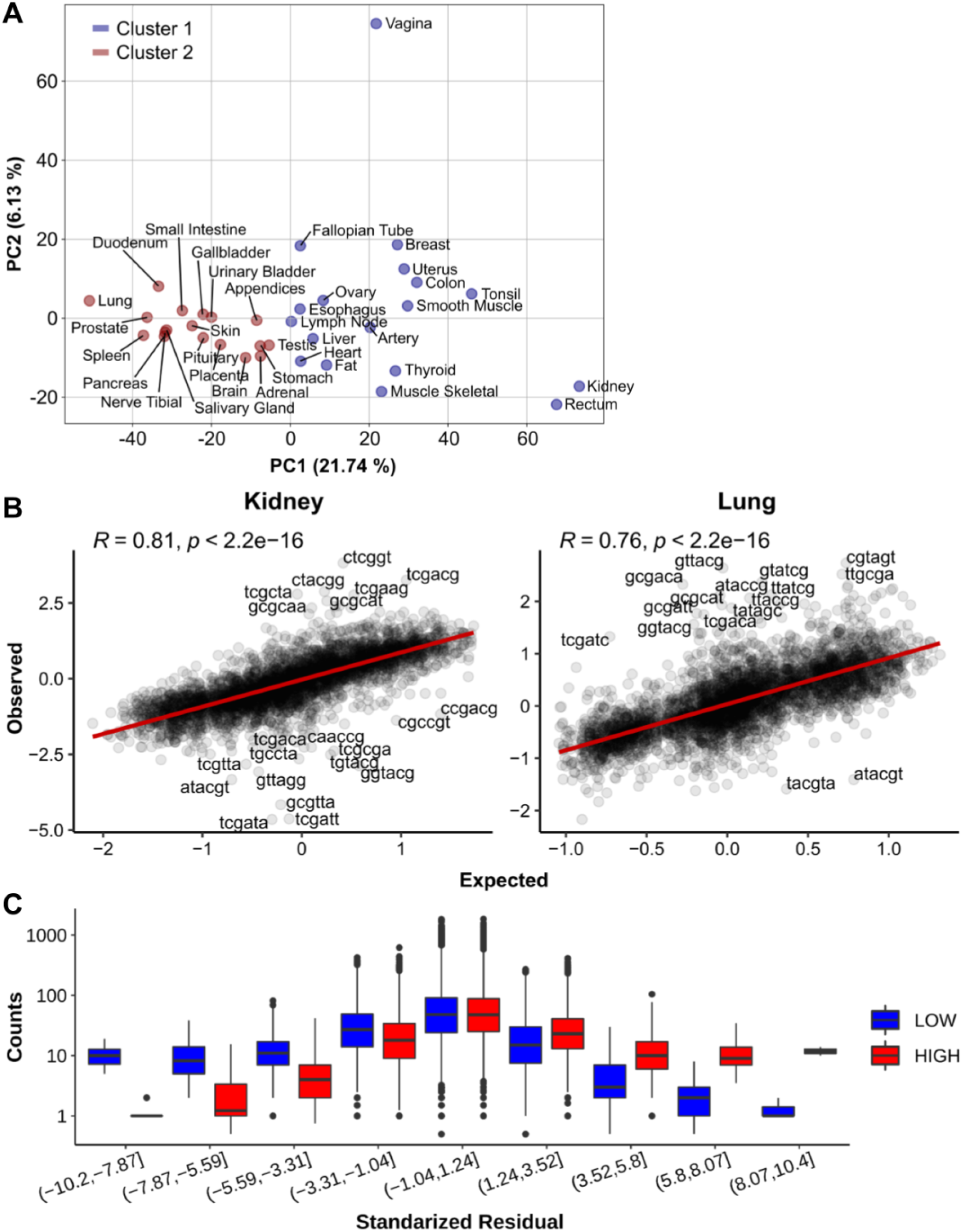
Differences between tissues are also observed at the codon pair level. (A) Principal Component Analysis (PCA) of the log-2-transformed ratios between high-PTR and low-PTR codon pair usage. The first axis completely separates the same two clusters detected in Fig. 2. (B) Linear regression in kidney and lung between the observed log-2-transformed codon pair ratios and the expected values based on the contribution of their constituent codons alone (see Methods). Codon pairs with a standardized residual higher than 4 are labeled. (C) For each standardized residual interval, number of codon pair counts in high-PTR and low-PTR gene sets. All tissues are merged together.

**Extended Data Figure 5.**
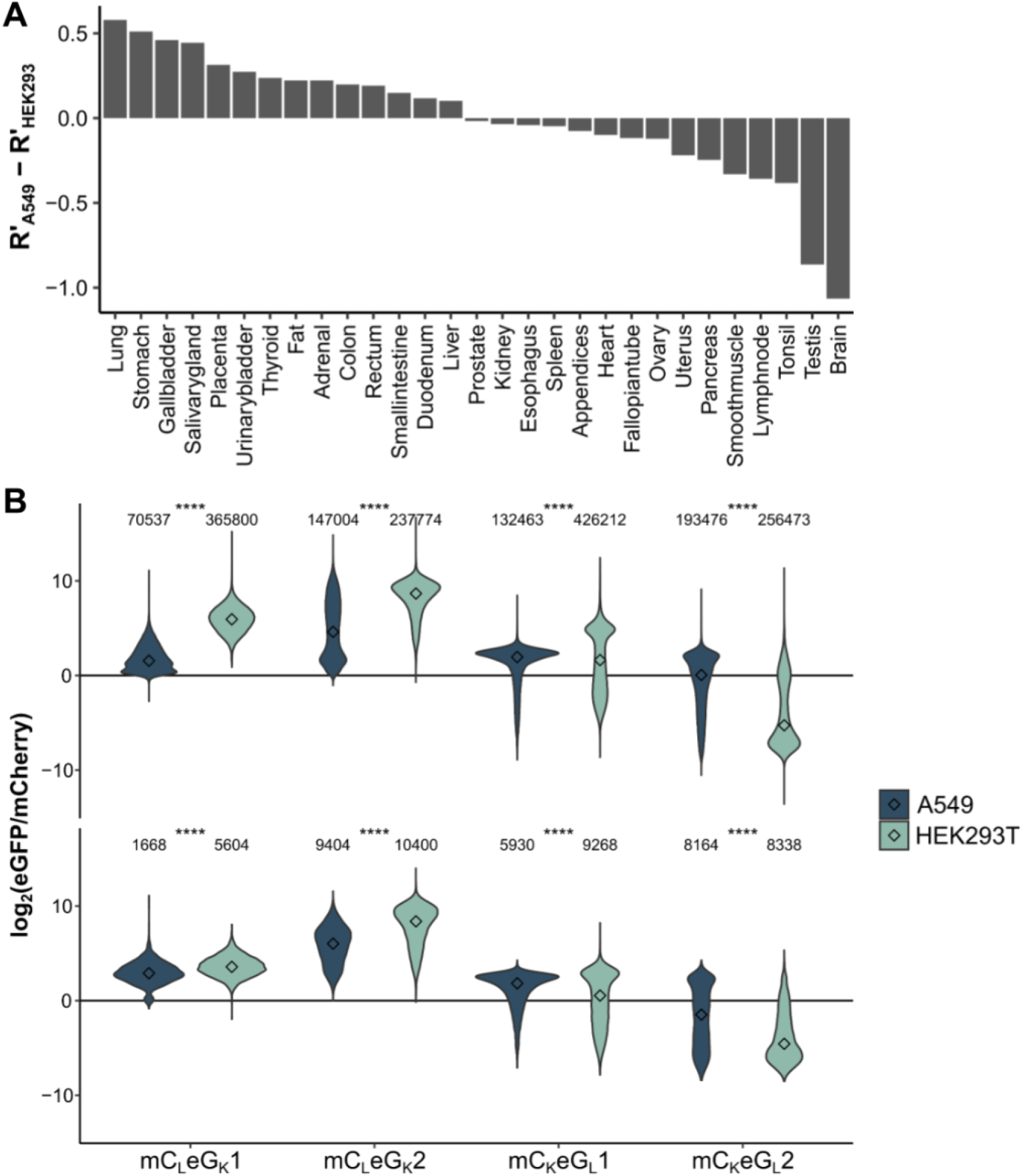
CUSTOM generates fluorescent variants with desired tissue-specific expression, related to Figure 3. (A) Difference between the standardized Spearman correlation of the proteomics profiles of A549 and HEK293^25^ against all tissues in the HPA^20^. (B) Ratios of eGFP and mCherry for each of the four constructs detected by flow cytometry. The number of cells within each group is specified. Center values represent the median. Statistical differences were determined by two-tailed Wilcoxon rank-sum test, and are denoted as follows: *p ≤ 0.05, **p ≤ 0.01, ***p ≤ 0.001, ****p ≤ 0.0001.

## SUPPLEMENTARY TABLES

**Supplementary Table 1.** Protein-to-mRNA ratios among tissues. (A) Protein-to-mRNA ratios of the Human Protein Atlas. (B) Protein-to-mRNA ratios of GTEx. (C) GTEx samples. (D) Translational efficiency of brain, liver and testis. (E) High-PTR and low-PTR sets of proteins for each tissue. (F) Concordance of protein sets between tissues in both HPA and GTEx. (G) Protein half-lives of five different human cell lines. (H) Proteins secreted to blood.

**Supplementary Table 2.** Random Forest models of codon usage. (A) Results of RF models for each of the 36 tissues. (B) Codon ratios between high-PTR vs low-PTR proteins for each tissue. (C) Codon ratios between high-PTR vs low-PTR proteins for each tissue, after excluding secretome proteins. (D) Random Forest models based on codon usage of misframed CDSs and dinucleotide usage.

**Supplementary Table 3.** Models of codon pair usage. (A) Codon pair ratios between high-PTR vs low-PTR proteins for each tissue. (B) Linear regression between observed vs expected codon pair ratios.

**Supplementary Table 4.** TisOpt-optimized fluorescent variants. (A) Relative Codon Usage of optimized protein variants. (B) Sequences of optimized protein variants.

## REFERENCES

1. Ranaghan, M. J., Li, J. J., Laprise, D. M. & Garvie, C. W. Assessing optimal: inequalities in codon optimization algorithms. BMC Biol. 19, 36 (2021).

2. Bergman, S. & Tuller, T. Widespread non-modular overlapping codes in the coding regions. Phys. Biol. 17, 031002 (2020).

3. Gould, N., Hendy, O. & Papamichail, D. Computational Tools and Algorithms for Designing Customized Synthetic Genes. Front. Bioeng. Biotechnol. 2, (2014).

4. Watts, A., Sankaranarayanan, S., Watts, A. & Raipuria, R. K. Optimizing protein expression in heterologous system: Strategies and tools. Meta Gene 29, 100899 (2021).

5. Tunney, R. et al. Accurate design of translational output by a neural network model of ribosome distribution. Nat. Struct. Mol. Biol. 25, 577–582 (2018).

6. Medina-Muñoz, S. G. et al. iCodon: ideal codon design for customized gene expression. bioRxiv 2021.05.06.442969 (2021) doi:10.1101/2021.05.06.442969.

7. Mordstein, C. et al. Codon Usage and Splicing Jointly Influence mRNA Localization. Cell Syst. 10, 351–362.e8 (2020).

8. Chen, S. et al. Codon-Resolution Analysis Reveals a Direct and Context-Dependent Impact of Individual Synonymous Mutations on mRNA Level. Mol. Biol. Evol. 34, 2944–2958 (2017).

9. Dittmar, K. A., Goodenbour, J. M. & Pan, T. Tissue-Specific Differences in Human Transfer RNA Expression. PLOS Genet. 2, e221 (2006).

10. Hernandez-Alias, X., Benisty, H., Schaefer, M. H. & Serrano, L. Translational efficiency across healthy and tumor tissues is proliferation-related. Mol. Syst. Biol. 16, e9275 (2020).

11. Kames, J. et al. TissueCoCoPUTs: Novel Human Tissue-Specific Codon and Codon-Pair Usage Tables Based on Differential Tissue Gene Expression. J. Mol. Biol. 432, 3369–3378 (2020).

12. Frumkin, I. et al. Codon usage of highly expressed genes affects proteome-wide translation efficiency. Proc. Natl. Acad. Sci. U. S. A. 115, E4940–E4949 (2018).

13. Wu, C. C.-C., Zinshteyn, B., Wehner, K. A. & Green, R. High-Resolution Ribosome Profiling Defines Discrete Ribosome Elongation States and Translational Regulation during Cellular Stress. Mol. Cell 73, 959–970.e5 (2019).

14. Buccitelli, C. & Selbach, M. mRNAs, proteins and the emerging principles of gene expression control. Nat. Rev. Genet. 21, 630–644 (2020).

15. GTEx Consortium. The Genotype-Tissue Expression (GTEx) project. Nat. Genet. 45, 580–585 (2013).

16. Uhlén, M. et al. Tissue-based map of the human proteome. Science 347, (2015).

17. Eraslan, B. et al. Quantification and discovery of sequence determinants of protein-per-mRNA amount in 29 human tissues. Mol. Syst. Biol. 15, e8513 (2019).

18. Franks, A., Airoldi, E. & Slavov, N. Post-transcriptional regulation across human tissues. PLOS Comput. Biol. 13, e1005535 (2017).

19. Jiang, L. et al. A Quantitative Proteome Map of the Human Body. Cell 183, 269–283.e19 (2020).

20. Wang, D. et al. A deep proteome and transcriptome abundance atlas of 29 healthy human tissues. Mol. Syst. Biol. 15, e8503 (2019).

21. Wang, Z.-Y. et al. Transcriptome and translatome co-evolution in mammals. Nature 588, 642–647 (2020).

22. Mathieson, T. et al. Systematic analysis of protein turnover in primary cells. Nat. Commun. 9, 689 (2018).

23. Zecha, J. et al. Peptide Level Turnover Measurements Enable the Study of Proteoform Dynamics *. Mol. Cell. Proteomics 17, 974–992 (2018).

24. Coleman, J. R. et al. Virus Attenuation by Genome-Scale Changes in Codon Pair Bias. Science 320, 1784–1787 (2008).

25. Geiger, T., Wehner, A., Schaab, C., Cox, J. & Mann, M. Comparative Proteomic Analysis of Eleven Common Cell Lines Reveals Ubiquitous but Varying Expression of Most Proteins *. Mol. Cell. Proteomics 11, (2012).

26. Kudla, G., Lipinski, L., Caffin, F., Helwak, A. & Zylicz, M. High Guanine and Cytosine Content Increases mRNA Levels in Mammalian Cells. PLOS Biol. 4, e180 (2006).

27. Hia, F. et al. Codon bias confers stability to human mRNAs. EMBO Rep. 20, e48220 (2019).

28. Allen, S. R. et al. Distinct responses to rare codons in select Drosophila tissues. bioRxiv 2022.01.06.475284 (2022) doi:10.1101/2022.01.06.475284.

29. Lähnemann, D. et al. Eleven grand challenges in single-cell data science. Genome Biol. 21, 31 (2020).

30. Mauro, V. P. Codon Optimization in the Production of Recombinant Biotherapeutics: Potential Risks and Considerations. BioDrugs 32, 69–81 (2018).

31. Alexaki, A. et al. Effects of codon optimization on coagulation factor IX translation and structure: Implications for protein and gene therapies. Sci. Rep. 9, 15449 (2019).

32. Lorenz, R. et al. ViennaRNA Package 2.0. Algorithms Mol. Biol. 6, 26 (2011).

33. Sharp, P. M. & Li, W.-H. The codon adaptation index-a measure of directional synonymous codon usage bias, and its potential applications. Nucleic Acids Res. 15, 1281–1295 (1987).

34. Wright, F. The ‘effective number of codons’ used in a gene. Gene 87, 23–29 (1990).

35. Alexaki, A. et al. Codon and Codon-Pair Usage Tables (CoCoPUTs): Facilitating Genetic Variation Analyses and Recombinant Gene Design. J. Mol. Biol. 431, 2434–2441 (2019).

36. Pujar, S. et al. Consensus coding sequence (CCDS) database: a standardized set of human and mouse protein-coding regions supported by expert curation. Nucleic Acids Res. 46, D221–D228 (2018).

37. Pedregosa, F. et al. Scikit-learn: Machine Learning in Python. J. Mach. Learn. Res. 12, 2825−2830 (2011).

38. Merico, D., Isserlin, R., Stueker, O., Emili, A. & Bader, G. D. Enrichment map: a network-based method for gene-set enrichment visualization and interpretation. PloS One 5, e13984 (2010).

39. Ashburner, M. et al. Gene Ontology: tool for the unification of biology. Nat. Genet. 25, 25–29 (2000).

40. Mi, H., Muruganujan, A., Ebert, D., Huang, X. & Thomas, P. D. PANTHER version 14: more genomes, a new PANTHER GO-slim and improvements in enrichment analysis tools. Nucleic Acids Res. 47, D419–D426 (2019).

41. Gremse, M. et al. The BRENDA Tissue Ontology (BTO): the first all-integrating ontology of all organisms for enzyme sources. Nucleic Acids Res. 39, D507–513 (2011).

